# RIPK1 activates distinct gasdermins in macrophages and neutrophils upon pathogen blockade of innate immune signalling

**DOI:** 10.1101/2021.01.20.427379

**Authors:** Kaiwen W. Chen, Benjamin Demarco, Rosalie Heilig, Saray P Ramos, James P Grayczyk, Charles-Antoine Assenmacher, Enrico Radaelli, Leonel D. Joannas, Jorge Henao-Mejia, Igor E Brodsky, Petr Broz

## Abstract

Injection of effector proteins to block host innate immune signalling is a common strategy used by many pathogenic organisms to establish an infection. Pathogenic *Yersinia* species for example inject the acetyltransferase YopJ into target cells to inhibit NF-κB and MAPK signalling. To counteract this, detection of YopJ activity in myeloid cells promotes the assembly of a RIPK1-caspase-8 death-inducing platform that confers antibacterial defence. While recent studies revealed that caspase-8 cleaves the pore-forming protein, gasdermin D (GSDMD) to trigger pyroptosis in macrophages, whether RIPK1 activates additional substrates downstream of caspase-8 to promote host defence is unclear. Here, we report that the related gasdermin family member gasdermin E (GSDME) is activated upon detection of YopJ activity in a RIPK1 kinase-dependent manner. Specifically, GSDME promotes neutrophil pyroptosis and IL-1β release, which is critical for anti-*Yersinia* defence. During *in vivo* infection, IL-1β neutralisation increases bacterial burden in wild type but not *Gsdme*-deficient mice. Thus, our study establishes GSDME as an important mediator that counteracts pathogen blockade of innate immune signalling.

## Introduction

Gasdermins are a family of recently described pore-forming proteins and are emerging as key drivers of cell death and inflammation. Gasdermins comprise of a cytotoxic N-terminal domain connected to an inhibitory C-terminal domain and are activated upon proteolytic cleavage (1, 2). This cleavage event releases the cytotoxic N-terminal fragment, which creates membrane pores and trigger a form of lytic cell death called pyroptosis. Gasdermin D (GSDMD) is arguably the best characterised family member to date, and is activated upon proteolysis by caspase-1, −4, −5, −8, −11 and serine proteases (3–10). Active GSDMD promotes host defence by eliminating the replicating niche of intracellular pathogens (11) and inducing the extrusion of antimicrobial neutrophil extracellular traps (NETs) (12). In addition, GSDMD pores act as a conduit for bioactive IL-1β release (13–15), a potent pro-inflammatory cytokine that similarly requires proteolytic cleavage by caspase-1 or −8 to gain biological activity (16). By contrast, GSMDE (also known as DFNA5) is activated by apoptotic caspase-3 and −7 and granzyme B, which drives tumour cell pyroptosis and anti-tumour immunity (17–19). The physiological function of GSDME in primary immune cells and its potential role in host defence remains unresolved and has not been reported.

Pathogenic *Yersinia* are a group of Gram-negative extracellular bacteria which cause disease ranging from gastroenteritis (*Y. pseudotuberculosis*) to plague (*Y. pestis*). A major mechanism by which pathogenic *Yersinia* establish systemic infection is by injecting the effector protein YopJ, an acetyltransferase that blocks transforming growth factor beta-activated kinase 1 (TAK1) to inhibit host innate immune signalling and proinflammatory cytokine production (20). To counteract this, detection of YopJ activity by myeloid cells induces the assembly of a cytoplasmic death-inducing complex that comprises of receptor-interacting serine/threonine protein kinase 1 (RIPK1), fas-associated protein with death domain (FADD) and caspase-8 (20–22). During *in vivo* infection, RIPK1/caspase-8-dependent cell death in myeloid cells restricts bacterial dissemination and replication at distal sites by inducing proinflammatory cytokine production from uninfected bystander cells (20). More recently, GSDMD was identified as a novel caspase-8 substrate during *Yersinia* infection that drives anti-microbial defence *in vivo* (7, 8, 23). However, whether RIPK1 activates additional substrates to restrict *Yersinia* infection is unclear and a focus of this study. Here, we identify GSDME as a novel substrate activated downstream of RIPK1 that confers host resistance against *Yersinia*. *Gsdme-*deficient mice failed to control bacterial replication in the spleen and liver and consequently are more susceptible to *Yersinia* infection than WT animals. Mechanistically, our data reveal that RIPK1 promotes caspase-3-dependent GSDME activation and IL-1β release in neutrophils but not macrophages. Neutralisation of IL-1β impaired bacterial clearance in wild type (WT) but not *Gsdme*^−/−^ animals, indicating that GSDME pores promote the major part of IL-1β release during *Yersinia* challenge *in vivo.*

### RIPK1 promotes GSDMD-dependent and independent antimicrobial response

We and others recently demonstrated that RIPK1-dependent GSDMD activation promotes macrophage pyroptosis and host defence against pathogenic *Yersinia* infection *in vivo* (7, 8, 23), however, whether RIPK1 activates other substrates to promote anti-*Yersinia* defence is unclear. To investigate this possibility, we challenged WT, *Gsdmd*^−/−^ and *Ripk1*^*D138N/D138N*^ ‘kinase-dead’ mice (hereafter referred as *Ripk1*^*KD*^) with *Yersinia pseudotuberculosis* (*Yptb*) and monitored their survival. In agreement with previous report (20), RIPK1 kinase activity is critical for anti-*Yersinia* defence, as *Ripk1*^*KD*^ mice exhibited 100% mortality within 7 days of infection, whereas more than 70% of WT animals survived 14 days post-infection **(Fig. 1A)**. Surprisingly, we observed that *Gsdmd*^-^deficiency does not fully recapitulate the susceptibility of *Ripk1*^*KD*^ mice to *Yptb* infection **(Fig. 1A)**, indicating that RIPK1 kinase activity activates additional unidentified substrates to confer anti-*Yersinia* defence. Since RIPK1 kinase activity promotes caspase-8-dependent apoptotic caspase activation (24), and apoptotic caspases-3/7 cleaves GSDME (17, 19), we investigated whether GSDME drives anti-bacterial defence. Indeed, *Gsdme*^−/−^ mice were significantly more susceptible to *Yptb* infection compared to WT controls **(Fig. 1B)** and significantly higher bacterial burden were recovered from the spleen and liver of *Gsdme*^−/−^ mice compared to WT controls **(Fig. 1C-D)**. In contrast, comparable bacteria were recovered from the mesenteric lymph nodes (MLN) and Peyer’s patches (PP) of WT and *Gsdme*^−/−^ mice **(Fig. 1E-F)**, indicating that GSDME primarily functions to prevent *Yptb* dissemination and replication at distal sites. These observations are consistent with the reported function of RIPK1 in myeloid cells during *Yersinia* infection (20).

**Figure 1.**
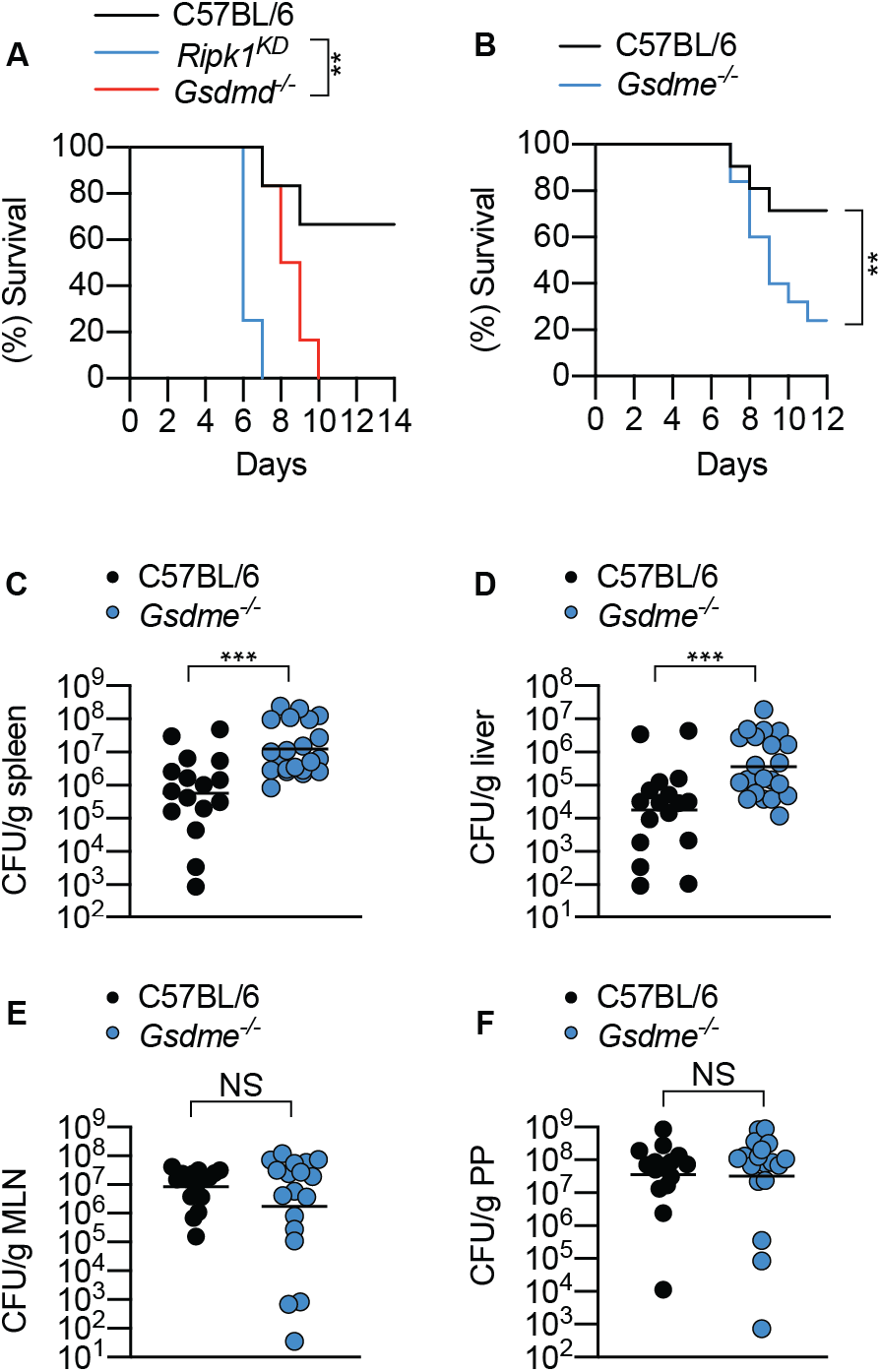
GSDME promotes host defence against *Y. pseudotuberculosis* infection. (A-F) Mice were challenged with 2 x 10^8^ CFU *Y. pseudotuberculosis.* (C-F) Bacterial load in the (C) spleen, (D) liver, (E) mesenteric lymph node and (F) Peyer’s patch were quantified at 5 days post infection. Data are geometric mean pooled from (C-F) 2 independent experiments. (E) Survival curve are (A) representative of two experiments or (B) pooled from 3 independent experiments. **P < 0.01 or ***P < 0.001.

### GSDME is dispensable for macrophage pyroptosis or cytokine secretion upon Y. pseudotuberculosis infection

Because RIPK1 kinase activity drives anti-*Yersinia* defence through the myeloid compartment (20), and *Yersinia* infection promotes RIPK1-dependent cell death in macrophages (7, 8, 20–22), we next focused our studies using bone marrow-derived macrophages (BMDMs). Consistent with previous reports (7, 8, 23), *Yptb* infection triggered robust processing of full-length GSDMD into its active p30 and inactive p20 fragment **(Fig. 2A)**, and corresponding GSDMD-dependent pyroptosis in WT macrophages **(Fig. 2B)**. Interestingly, while GSDME was completely processed into its active p30 fragment **(Fig. 2A)**, LDH release between WT and *Gsdme*^−/−^ BMDMs was indistinguishable **(Fig. 2B)**. *Gsdme-*deficiency did not further reduce LDH release in *Gsdmd*^−/−^ macrophages **(Fig. 2B)**, indicating that GSDME is dispensable for macrophage pyroptosis in both WT and *Gsdmd*^−/−^ macrophages during *Yptb* infection. Gasdermin-dependent pyroptosis is often tightly coupled with the release of mature IL-1β. In keeping with this, we observed that GSDMD but not GSDME is required for IL-1β secretion from *Yptb-*infected BMDMs **(Fig. 2C-D)**. Collectively, these data indicate that while GSDME is processed into its p30 active fragment during *Yptb* infection, is dispensable for macrophage pyroptosis and cytokine secretion.

**Figure 2.**
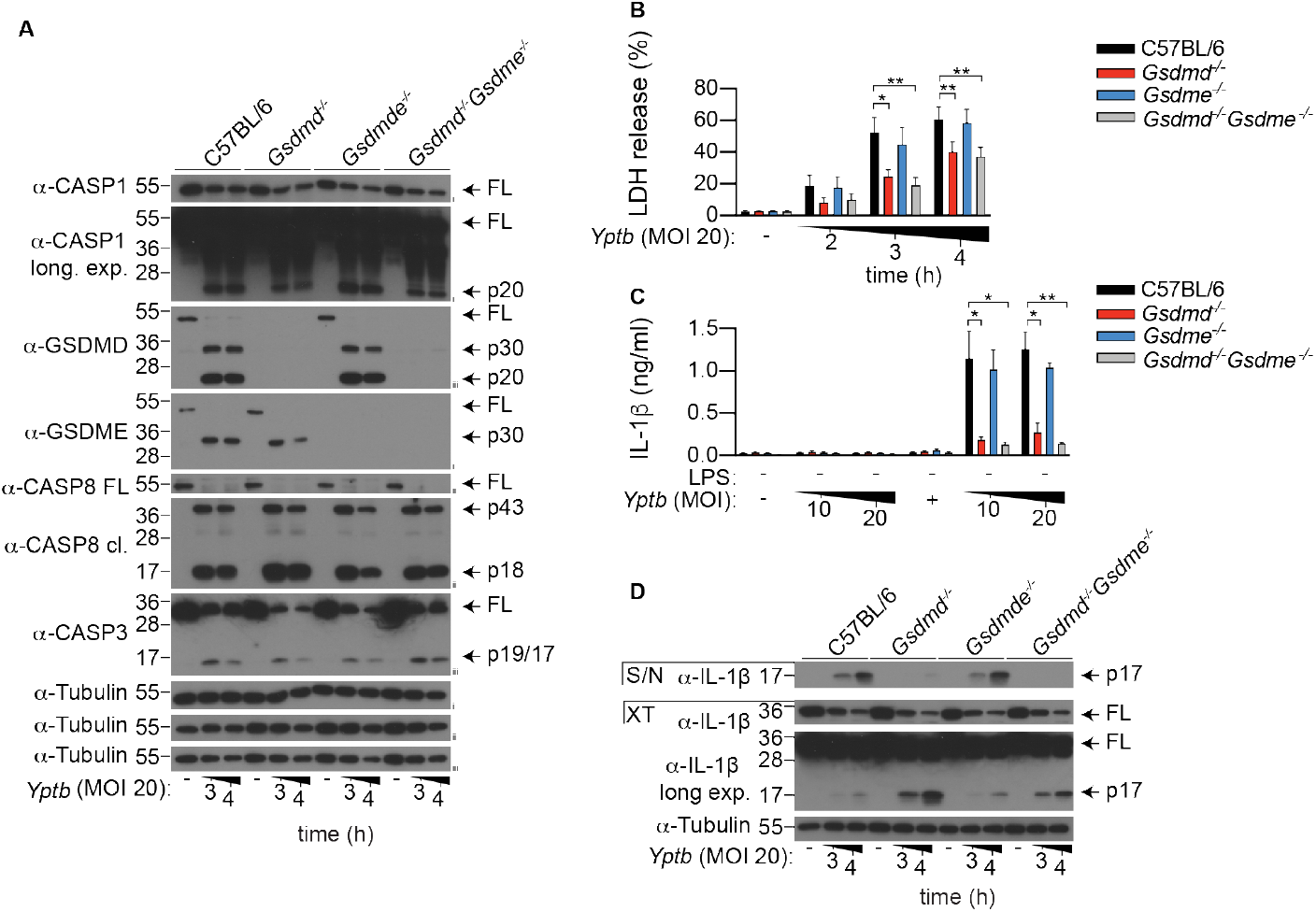
GSDME is dispensable for pyroptosis and cytokine secretion upon *Y. pseudotuberculosis* infection in macrophages. (A-B) Unprimed or (C-D) LPS-primed BMDMs were infected with *Y. pseudotuberculosis* (*Yptb*) for the (A-B, D) indicated time points or (C) for 3 h. (A) Mixed supernatant and cell extracts were examined by immunoblotting. (A-D) Immunoblots are representative of 3 independent experiments. (B-C) Data are mean + SEM of pooled data from 4 independent experiments. *P < 0.05, **P < 0.01.

### GSDME but not GSDMD promotes apoptotic neutrophil cell lysis

Neutrophils play an important role in host defence against *Yersinia* infection (25, 26) and are the most abundant cell type that are injected with effector proteins during *Yersinia* infection (27). Therefore, we next examined the contribution of GSDMD and GSDME to neutrophil cell lysis and cytokine secretion following *Yptb* infection. Under *in vitro* conditions, unstimulated neutrophils undergo spontaneous activation of the intrinsic apoptosis pathway and progress into lytic cell death through an ill-defined mechanism (28). Surprisingly, while purifying bone marrow neutrophils for *Yersinia* infection, we observed that naïve *Gsdme*^−/−^ neutrophils were significantly protected from spontaneous cell lysis, as uptake of the membrane-impermeable nucleic acid dye, propidium iodide (PI), was significantly reduced in *Gsdme*^−/−^ neutrophils compared to WT controls **(Fig. 3A)**. By contrast, PI uptake between WT and *Gsdmd*^−/−^ neutrophils was indistinguishable **(Fig. 3A)**, consistent with previous reports (3, 12, 15, 29). *Gsdmd/Gsdme double-*deficiency did not further reduce PI uptake compared to *Gsdme*^−/−^ neutrophils **(Fig. 3A)**. The pan-caspase inhibitor, Q-VD-Oph (QVD), supressed caspase-3 and GSDME activation in unstimulated neutrophils **(Fig. 3B)** and triggered a corresponding reduction in PI uptake compared to unstimulated WT and *Gsdmd*^−/−^ neutrophils **(Fig. 3C)**. QVD did not further reduce PI uptake in *Gsdme*^−/−^ and *Gsdmd*^−/−^*Gsdme*^−/−^ neutrophils compared to unstimulated cells **(Fig. 3D)**. In line with a previous report (17), we did not detect any difference in neutrophil numbers between naïve WT and *Gsdme*^−/−^ animals *in vivo* **(Fig. S1A-B)**. Collectively, these data demonstrate that GSDME is the major driver of spontaneous neutrophil lysis *in vitro* but does not regulate neutrophil turnover in naïve animals, most likely since *Gsdme*-deficiency does not block apoptosis of neutrophils but rather re-routes it to a different outcome.

**Figure 3.**
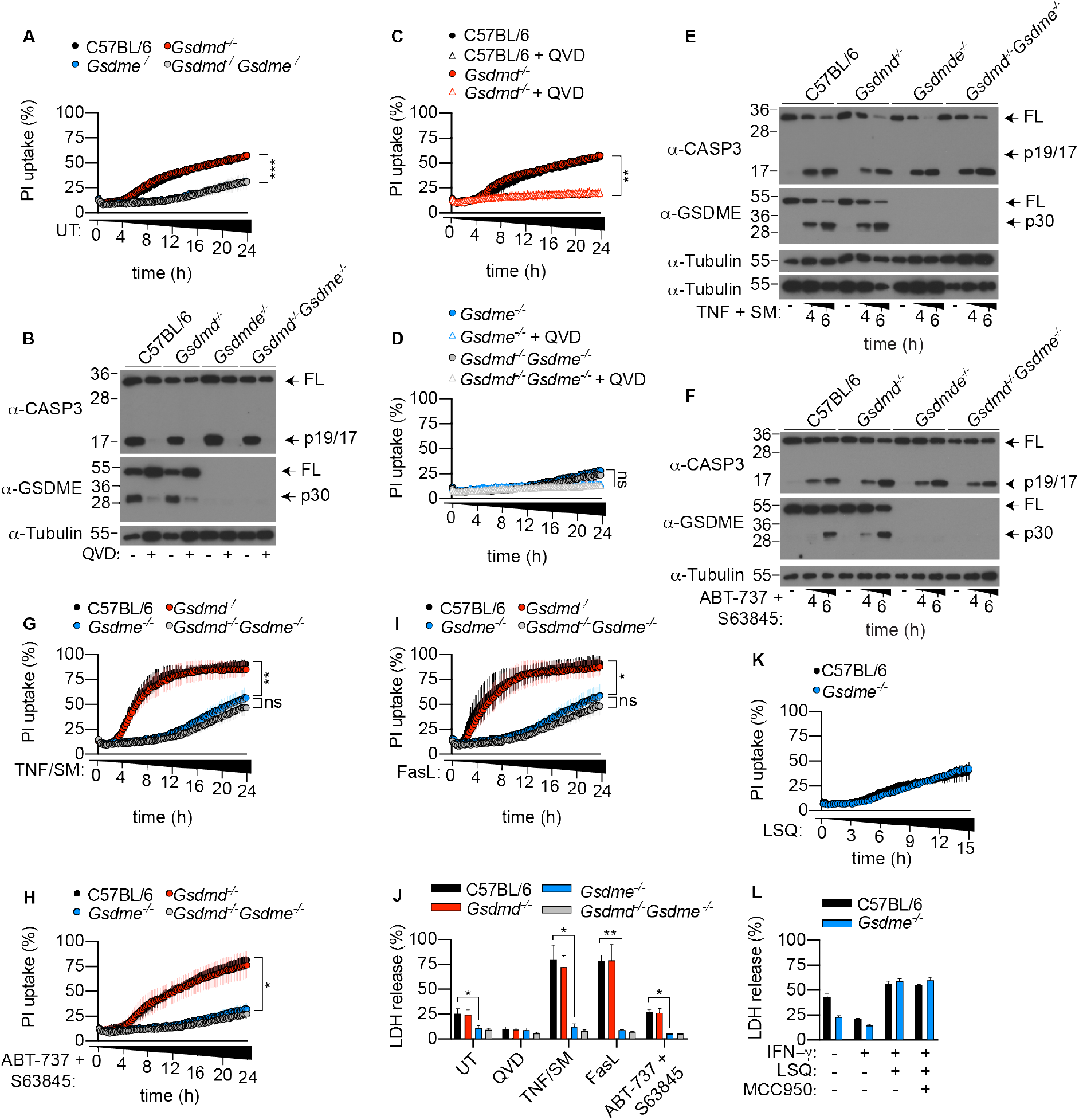
GSDME activation is required for neutrophil lysis upon apoptotic caspase activation. (A, C-D, G-I, K) Propidium iodine (PI) uptake was quantified over 24 h. (B, E, F) Mixed supernatant and cell extracts were examined by immunoblotting, representative of 3 independent experiments. (J-K) LDH release was measured at (J) 6 or (L) 15 h post-stimulation. Data are mean + SEM pooled from 3 (A, C, D, H, J) or 4 (G, I) independent experiments. (K-L) Data are mean + SD for technical triplicates representative from 3 independent experiments. *P < 0.05, **P < 0.01, ***P<0.001.

We next investigated whether GSDME drives neutrophil lysis following activation of extrinsic and intrinsic apoptosis. For this, we exposed neutrophils to TNF plus SMAC mimetic (SM) or FasL to induce extrinsic apoptosis, or ABT-737 plus S63845 to induce intrinsic apoptosis. Both extrinsic and intrinsic apoptosis triggered robust caspase-3 and GSDME activation **(Fig. 3E-F)**, and corresponding GSDME-dependent cell permeability and lysis, as measured by PI uptake and LDH release respectively **(Fig. 3G-J)**. In contrast, *Gsdmd-*deficiency did not impact PI uptake and LDH release in WT or *Gsdme*^−/−^ neutrophils **(Fig. 3G-J)**. Next, we stimulated neutrophils with LPS, SM and QVD (LSQ) to induce RIPK3-dependent necroptosis **(Fig. S2A-B)** (30). As anticipated, PI uptake and LDH release between WT and *Gsdme*^−/−^ neutrophils were comparable after LSQ stimulation **(Fig. 3K-L)**, indicating that *Gsdme*^−/−^ neutrophils are not intrinsically resistant to cell lysis. Collectively, this indicates that GSDME but not GSDMD promotes cellular lysis in apoptotic neutrophils.

### GSDME drives neutrophil pyroptosis and cytokine secretion upon Yersinia infection

We next sought to characterise the role of neutrophil GSDMD and GSDME during *Yptb* infection. We first primed neutrophils with IFN-γ in order to reduce spontaneous cell lysis (31) **(Fig. S3A-B)** and LPS to induce pro-IL-1β expression, prior to *Yptb* infection. Indeed, *Yptb* infection triggered GSDME-dependent neutrophil pyroptosis **(Fig. 4A)** and corresponding GSDME cleavage **(Fig. 4B)**, which is dependent on the bacterial effector YopJ and the RIPK1 kinase activity **(Fig. S3C)**. By contrast, GSDMD was dispensable on its own, and did not further contribute to lysis in the absence of GSDME **(Fig. 4A)**. Since GSDMD activation does not universally trigger pyroptosis in neutrophils (12, 29, 32, 33), we next investigated the cleavage status of neutrophil GSDMD upon *Yptb* infection. WT and *Caspase-1/11*^−/−^ neutrophils were challenged with *Yptb* or a *Salmonella ΔsifA* mutant, which triggers robust caspase-11-dependent GSDMD cleavage, as a positive control (12). In keeping with previous reports (12, 33), we observed robust processing of full length GSDMD into the active p30 fragment upon caspase-11 activation in WT neutrophils **(Fig. 4C)**. By contrast, GSDMD processing into the active p30 fragment was significantly weaker upon *Yptb* infection, while the inactive p43 and p20 species were more abundant **(Fig. 4C)** (4, 34). This suggests that in neutrophils, apoptotic caspases inactivate GSDMD during *Yptb* infection. In keeping with macrophage studies (7, 8), GSDMD processing was sensitive to the RIPK1 kinase inhibitor Nec-1s, and only partially reduced in *Caspase-1/11*^−/−^ neutrophils compared to WT controls upon *Yptb* infection **(Fig. 4C)**. These data demonstrate that while both caspase-1 and −8 promote GSDMD processing; neutrophil pyroptosis upon *Yptb* infection is solely driven by GSDME.

**Figure 4.**
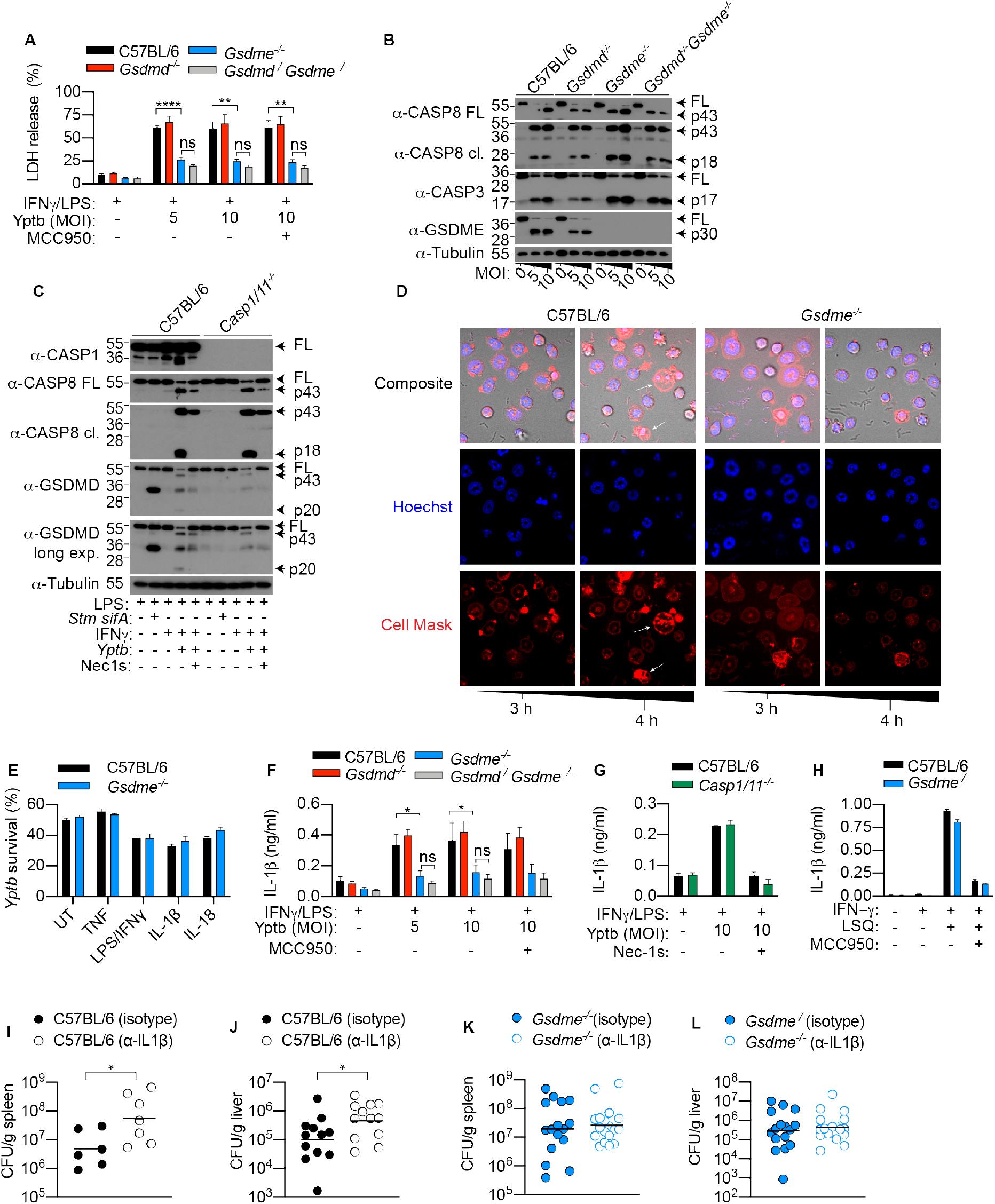
GSDME-dependent IL-1β promotes anti-*Yersinia* defence. (A-D, F-G) Neutrophils were infected with *Y. pseudotuberculosis* (*Yptb*) or *Salmonella ΔsifA* (both MOI 10) for 4 hours. (B, C) Mixed supernatant and cell extracts were examined by immunoblotting, representative of 3 independent experiments. (D) Time-lapse confocal images of *Yptb*-infected neutrophils. (E) Neutrophils were left unstimulated or primed with the indicated cytokines or PAMPs for 2 h prior to *Yptb* (MOI 1) infection for 3 h in antibiotic-free media. *Yptb* survival relative to time inoculum were displayed. (H) Neutrophils were treated with LPS/Smac mimetic/QVD (LSQ) for 15 h. (I-K) Mice were administered with isotype control or α-IL-1β neutralizing and challenged with *Y. pseudotuberculosis* for 5 days. (A, F) Data are mean + SEM pooled from 4 independent experiments, (I-L) geometric mean of pooled from 2 independent experiments or (E, G, H) + SD of triplicate stimulation representative of 3 independent experiments. *P < 0.05, **P < 0.01, **8*P<0.0001.

Since *Yptb* is an extracellular bacterium, and caspase-11-driven activation of GSDMD promotes the release of antimicrobial neutrophil extracellular traps (NETs) (12), we examined whether GSDME activation also triggers NET extrusion to restrict *Yptb* replication *in vitro*. Interestingly, while *Yptb* infection triggered robust GSDME processing and pyroptosis **(Fig. 4A-B)**, it did not result in any hallmarks of NETosis including nuclear decondensation and delobuation, or the appearance of diffused or spread NETs **(Fig. 4D)** (10). Instead, *Yptb* infection triggered morphological hallmarks of pyroptosis including membrane ballooning and nuclear condensation in a GSDME-dependent manner **(Fig. 4D)**, indicating that GSDME activation does not promote NET extrusion. In support of this, WT and *Gsdme*^−/−^ neutrophils killed *Yptb* to the same extent *in vitro* **(Fig. 4E, Fig. S4A-D)**. Since GSDME activation does not appear to directly restrict bacterial growth; and given that apoptotic neutrophils secrete the leaderless cytokine IL-1β (30, 35), we investigated whether GSDME activation drives the release of leaderless cytokines from pyroptotic neutrophils. In keeping with the cell lysis data **(Fig. 4A)**, IL-1β release from *Yptb-*infected neutrophils was indeed GSDME-dependent but GSDMD-independent **(Fig. 4F)**. Interestingly, while *Yptb* infection triggered caspase-1-dependent IL-1β maturation in macrophages (23), application of the NLRP3-specific inhibitor, MCC950, or *caspase-1/11*-deficiency, had no impact on GSDME-dependent IL-1β secretion from neutrophils **(Fig. 3F-G)**. This indicates that IL-1β maturation is largely caspase-8-dependent in apoptotic neutrophils, consistent with a previous report (30). In contrast, other leaderless IL-1 family cytokines including IL-1α and IL-33 were barely detected in the supernatant of pyroptotic neutrophils **(Fig. S5A-B)**. To ensure that the observed defect in IL-1β secretion is not due to a general secretion defect by *Gsdme*^−/−^ neutrophils, we stimulated IFNγ-primed neutrophils with a combination of LSQ to induce necroptosis and subsequent NLRP3-caspase-1 activation (30, 36). As anticipated, IL-1β secretion from necroptotic neutrophils is sensitive to the NLRP3-specific inhibitor, MCC950, and occurred independently of GSDME **(Fig. 4H)**, confirming that *Gsdme*^−/−^ neutrophils are not intrinsically defective in IL-1β release. Since neutrophils are a major cellular source of IL-1β during various bacterial infection (32, 37, 38), we investigated whether IL-1β release is a major mechanism by which GSDME promotes host resistance to *Yptb.* Indeed, IL-1β neutralisation triggered a significant increase in bacterial burden in the spleen and liver of WT animals compared to isotype control **(Fig. 4I-J)**, while the same regimen had no impact on bacterial clearance in *Gsdme*^−/−^ animals **(Fig. 4K-L)**. Collectively, these data observations reveal an unexpected role of GSDME as a major cellular driver of IL-1β release during *Yersinia* infection.

## Discussion

The discovery of gasdermin protein family has greatly revolutionised our understanding of cell death during microbial infection. However, majority of these studies focused solely on the archetypal gasdermin, GSDMD, for which a host of new and exciting functions including cytokine secretion (13–15), NET extrusion (10, 12) and repression of cGAS signalling (39) were described. By contrast, the physiological function of GSDME during microbial infection is unclear. In this study, we report that GSDME is a potent antimicrobial effector that defend against blockade of innate immune signalling by the Gram-negative bacteria *Y. pseudotuberculosis*. Although *Yersinia* infection triggered complete processing of full length GSDME into its active p30 fragment in macrophages, we found no evidence that GSDME promotes pyroptosis or IL-1β secretion in this cell type. This makes intuitive sense, as we recently reported that caspase-3/7-dependent GSDMD inactivation is required for optimal anti-*Yersinia* defence (23). Instead, we demonstrate that GSDME exerts its antimicrobial function in neutrophils by driving cellular pyroptosis and bioactive IL-1β release from YopJ-injected neutrophils. Although neutrophils were once regarded as effector cells that contribute minimally to orchestrate the immune response, this view is rapidly evolving as an increasing number of studies now document neutrophils as a major cellular source of proinflammatory cytokines (e.g., IL-1β and IL-8) during microbial infection (32, 37, 38, 40). Our study agrees with such a model, as we found that neutralisation of IL-1β impaired bacterial clearance in WT but not *Gsdme*^−/−^ animals, indicating that GSDME pores, and likely neutrophils, are a major cellular source of IL-1β during *Yersinia* infection. Since RIPK1 kinase activity in myeloid cells drives anti-*Yersinia* defence (20), it is possible that GSDME may also promote pyroptosis and bioactive IL-1β release from infected inflammatory monocytes. Given that gasdermin cleavage elicits distinct functional outcomes in different myeloid cell subsets (12, 29, 32, 33), future studies characterising the regulation and biological function of gasdermins in monocytes will be of interest. One potential caveat of our study, however, is the sensitivity between murine and human myeloid cells to *Yersinia* infection. While pathogenic *Yersinia* are often used to study cell death and gasdermin function in murine systems, human macrophages and neutrophils appear to be much more resistant to YopJ or small molecule TAK1 inhibitor-induced cell death compared to their murine counterparts (8, 41). This difference likely reflects higher expression of pro-survival molecules such as FLIP in human myeloid cells (42).

Our discovery that GSDME promotes neutrophil lysis and bioactive IL-1β release has major implications beyond infectious disease, as these processes are implicated in a variety of human diseases. For example, aberrant IL-1β secretion is associated with atherosclerosis, diabetes, neurological diseases, gout and rheumatoid arthritis. Because IL-1β secretion is significantly reduced in *Gsdmd-*deficient macrophages, GSDMD inhibitors are now regarded as attractive therapeutical targets for IL-1β-mediated disease. Our finding that GSDME promotes mature IL-1β release during *in vivo* infection unravels a previously unappreciated role for other gasdermins in driving IL-1β secretion. Future studies investigating the contribution of GSDME to such diseases may uncover novel therapeutics.

## Methods

### Animals

All experiments involving animals were performed under the guidelines and approval from the Swiss animal protection law (license VD3257) and guidelines from the National Institutes of Health (NIH) and the University of Pennsylvania Institutional Animal Use and Care Committee (Protocol #804523). C57BL/6J, *Ripk1*^*D138N/D138N*^ *Gsdmd*^−/−^, *Gsdme*^−/−^, *Gsdmd*^−/−^*Gsdme*^−/−^ and *Casp1/11*^*−/−*^ (all either generated in or back-crossed to the C57BL/6 background) were previously described (4, 43) and housed in specific-pathogen free facilities at the University of Lausanne. An independent line of *Gsdme*^−/−^ was generated at the University of Pennsylvania. *Gsdme* was knocked out in the C57BL/6J background by targeting exons 2 and 3 using two CRISPR gRNAs (CAGAACCCTCCTGTCACGAT and TGTGATATGGAGTACCCCGA) as previously described (44). Briefly, eggs were microinjected with the two gRNAs along with Cas9 mRNA. Progeny were screened for successful deletion and successfully deleted males were bred to C57BL/6J females. Germ-line transmitted founders were intercrossed to establish the *Gsdme*^−/−^ line.

### Primary myeloid cell culture

Bone marrow-derived macrophages were differentiated in DMEM (Gibco) containing 20% 3T3 supernatant (as a source of M-CSF), 10% heat-inactivated FCS (Bioconcept), 10 mM HEPES (Bioconcept), penicillin/streptomycin (Bioconcept) and non-essential amino acids (Gibco), and stimulated on day 7-9 of differentiation. Mature neutrophils were purified from murine bone marrow using anti-Ly6G-FITC (1A8 clone) and anti-FITC beads (Miltenyi) (>98% purity) as previously described (32). In all experiments, neutrophils were seeded at a density of 4×10^5^ cells per well in 200 µl Opti-MEM and stimulated on the day of purification, and macrophages were seeded at a density of 5×10^4^ cells per well in complete media a day prior to stimulation.

### Apoptosis and necroptosis assay

To activate extrinsic apoptosis, neutrophils were stimulated with recombinant murine TNF (100 ng/ml; Peprotech) and the SMAC mimetic AZD 5582 (0.25 μM; Selleckchem) or 100 ng/ml Fc-Fas (kind gift from Prof Pascal Schneider; University of Lausanne). To activate intrinsic apoptosis, neutrophils were stimulated with ABT-737 (500 nM; Selleckchem) and S63845 (500 nM; Selleckchem). To induce neutrophil necroptosis, cells were primed with recombinant murine IFN-γ (100 ng/ml; Peprotech) for 1 hour, followed by 2 hours incubation with *E. coli* 055:B5 LPS (100 ng/ml; Invivogen). Cells were treated with 10 μM Q-VD-Oph in the last 20-30 min of LPS priming, and stimulated with the SMAC mimetic AZD 5582 (0.25 μM; Selleckchem). Where indicated, neutrophils were treated with the NLRP3-specific inhibitor, MCC950 (10 μM; Invivogen) 20 minutes before SMAC mimetic treatment.

### *Yersinia* infection

Where indicated, macrophages were primed with *E. coli* 055:B5 LPS (100 ng/ml; Invivogen) for 3 hours prior to infection. For neutrophil infection, cells were co-primed with IFN-γ (100 ng/ml; Peprotech) and *E. coli* 055:B5 LPS (100 ng/ml; Invivogen) for 3 hours prior to infection. Log-phase *Yersinia pseudotuberculosis* strain 32777 were prepared and used to infect BMDM or neutrophils at a multiplicity of infection (MOI) 5-20 as previously described (23). IL-1α, IL-1β and IL-33 levels in cell-free supernatant were measured by ELISA according to manufacturers’ instruction (all R&D Systems DuoSet ELISA). For *in vivo* infection, mice were fasted for 16 h and challenged with 2 x 10^8^ CFU stationary phase bacteria by oral gavage. To determine bacterial burden, mice were euthanised 4- or 5-days post-infection and tissues were harvest, homogenised in 1 ml of PBS, and serially diluted on LB agar.

### *Salmonella* infection

Mature neutrophils were seeded at a density of 4×10^5^ cells per well in 200 μl Opti-MEM and primed with *E. coli* 055:B5 LPS (100 ng/ml; Invivogen) for 3 hours. *Salmonella enterica* serovar Typhimurium SL1344 *ΔsifA* mutant were grown overnight with aeration at 37 °C in LB media and stationary-phase bacteria were used to infect neutrophils as previously described (12).

### Cell death measurements

Cell permeabilization was quantified by measuring propidium iodide (1 μg/ml; Thermo Fisher Scientific) uptake over time using a fluorescent plate reader (Cytation5; Biotek). Cell lysis was quantified by measuring the amount of intracellular LDH release into the cell culture supernatant (TaKaRa LDH cytotoxicity detection kit; Clontech). Percentage PI uptake and LDH release were calculated relative to 100 % cell lysis in untreated control sample.

### Immunoblotting

Cell culture supernatant were precipitated with methanol and chloroform using standard methods and and resuspended in cell extracts lysed in boiling lysis buffer (66 mM Tris-Cl pH 7.4, 2% SDS, 10mM DTT, NuPage LDS sample buffer; Thermo Fischer). Proteins were separated on 14% polyacrylamide gels and transferred onto nitrocellulose membrane using Trans-blot Turbo (Bio-Rad). Antibodies for immunoblot were against GSDME (EPR19859; Abcam; 1:1,000), GSDMD (EPR19828; Abcam; 1:3000), full-length caspase-8 (4927; Cell Signaling; 1:1000), cleaved caspase-8 (9429; Cell Signalling; 1:1000), caspase-3 (9662; Cell Signalling; 1:1000), caspase-1 p20 (casper-1; Adipogen; 1:1000), pro-IL-1β (AF-401-NA; R&D; 1:1000) and alpha-tubulin (DM1A; Abcam; 1:5000).

### Flow cytometry

Bone marrow cells were blocked with TruStain FcX™ (anti-mouse CD16/32; Biolegend) and stained with CD11b (M1/70; Biolegend) and Ly6G (1A8; Biolegend) to identify neutrophil population. Blood neutrophils were identified as CD45.2^+^ (104; Biolegend) Gr-1^+^ (RB6-8C5; Biolegend) SSC^high^ cells. Cell profiles were acquired using BD Accuri C6 Plus or Cytek Aurora and analysed using FlowJo (Tree Star).

### Statistical analyses

Statistical analyses were performed using Prism 8 (Graphpad) software. Parametric *t* test was used for normally distributed data sets while non-normally distributed data sets were analyzed using non-parametric Mann-Whitney *t* tests. For PI uptake, area under the curve (AUC) was calculated for each sample and pooled data from 3-4 independent experiments were analyzed using a one-way ANOVA. Survival curves were compared using log-rank Mantel–Cox test. Data were considered significant when P ≤ 0.05, with *P ≤ 0.05 **P ≤ 0.01 or ***P ≤ 0.001.

## Acknowledgement

We thank Prof Manolis Pasparakis (University of Cologne, Germany) and Prof Pascal Schneider (University of Lausanne) for generously providing *Ripk1*^*D138N/D138N*^ mice and Fc-FasL respectively. We thank Dr Romain Bedel and Dr Anne Wilson (University of Lausanne) for assistance on flow cytometry and Dr Lance Peterson (Washington University in St Louis) for discussion. This work was supported by a European Research Council Grant (ERC2017-CoG-770988-InflamCellDeath) and a Swiss National Science Foundation project grant (310030_175576) to P.B., a National University of Singapore Start Up Grant and a Ministry of Education Inauguration Grant to K.W.C, and a Burroughs Wellcome Fund Pathogenesis of Infectious Disease Award to I.E.B.

## Author contributions

K.W.C., B.D., R.H., S.R.P., P.G., C-A.A. and E.R. performed experiments. L.D.J. and J.H-M. generated *Gsdme*^−/−^mice. I.E.B provided essential reagents and expert advice. K.W.C. and P.B. conceptualised and supervised the study and wrote the manuscript, which all authors reviewed before submission.

