## Supplementary material for "RIPK1 activates distinct gasdermins in macrophages and neutrophils upon pathogen blockade of innate immune signalling"

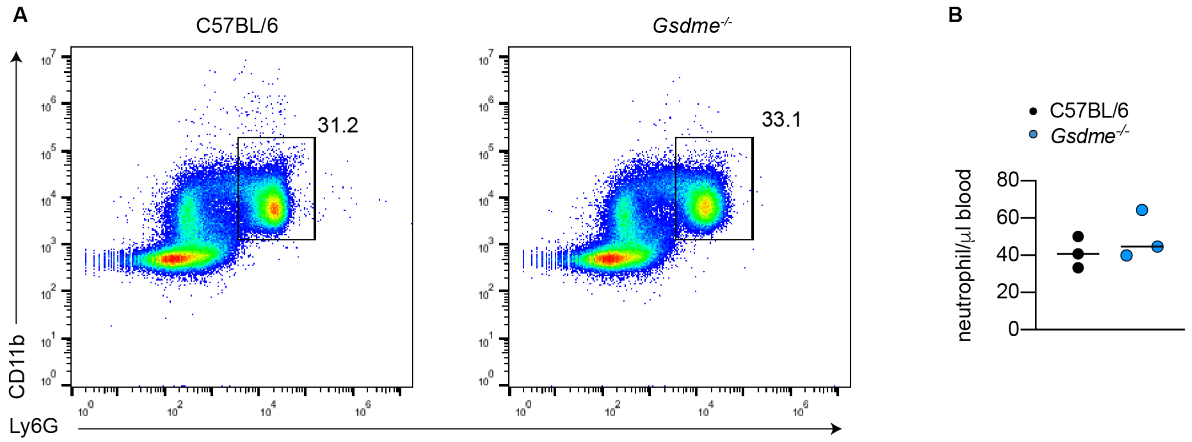


**Figure S1. GSDME does not regulate neutrophils numbers *in vivo.*** (A) Total bone marrow from 16-week old male WT and *Gsdme*$^{-/-}$ littermate controls were stained with CD11b and Ly6G and analysed by flow cytometry. Data are representative from 4 animals. (B) Blood neutrophils were quantified from 8-week old WT and *Gsdme*$^{-/-}$ mice by flow cytometry.

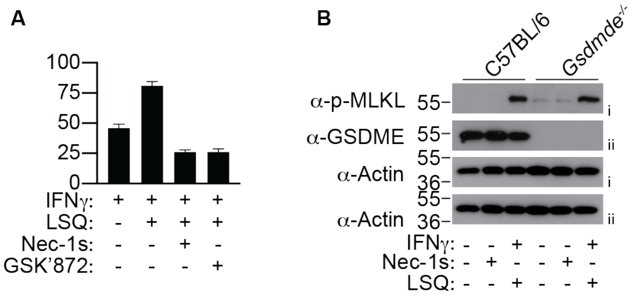


**Figure S2. LSQ triggers necroptosis in neutrophils.** (A) IFNγ-primed neutrophils were stimulated with LPS/SMAC mimetic/QVD (LSQ) for 16 h in the presence or absence of Nec-1s or GSK'872. (B) WT and and *Gsdme*$^{-/-}$ neutrophils were primed with IFNγ and stimulated with LSQ for 15 h and mixed supernantant and cell extracts were analysed by western blot. (A) Data are + SD from triplicate well stimulation representative of 2 experiments.

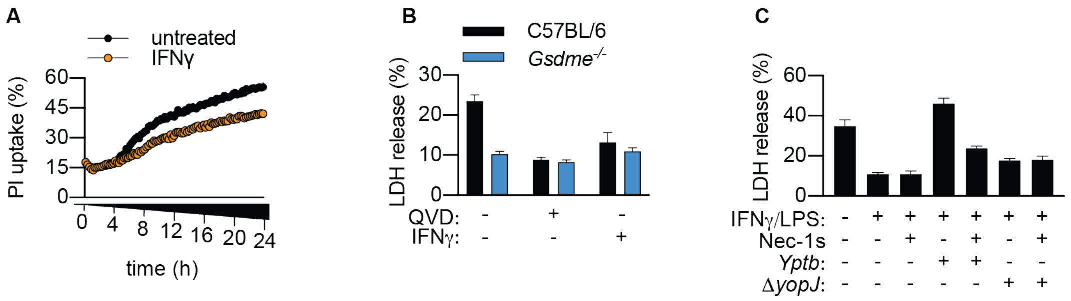


**Figure S3. IFNγ priming reduces spontaneous neutrophil lysis.** (A) Neutrophils were left untreated or stimulated with IFNγ and Propidium iodine (PI) uptake was quantified over 24 h. (B) Neutrophils were left untreated or stimulated with QVD or IFNγ and LDH release were quantified after 6 h. (C) Neutrophils were infection with *Yptb* or the *ΔyopJ* and LDH was quantified at 6 h. (A-C) Data are + SD from triplicate well stimulation representative of 2 experiments.

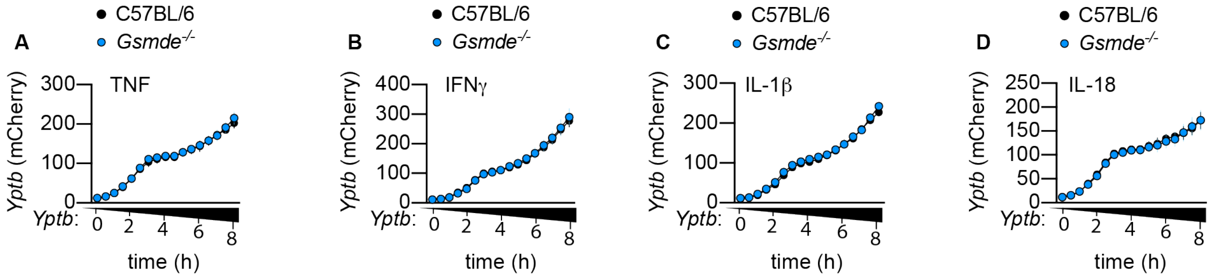


**Figure S4. GSDME does not promote bacterial clearance in a cell-intrinsic manner.** WT and *Gsdme*$^{-/-}$ neutrophils were primed with pro-inflammatory cytokines (all 100 ng/ml) for 3 h and challenged with *Y. pseudotuberculosis* (*Yptb*) for 8 h (MOI 1) and bacterial replication (mCherry) was quantified over time. (A-D) Data are mean + SD from triplicate well stimulation representative of 3 independent experiments.

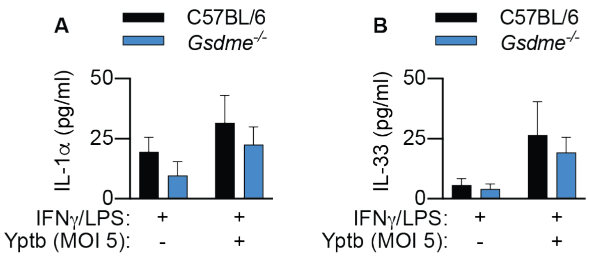


**Figure S5. Pyroptotic neutrophils do not release IL-1α or IL-33.** IFNγ/LPS-primed neutrophils were infected with *Y. pseudotuberculosis* (*Yptb*) for 4 hours. (A-B) Data are mean + SD for technical triplicates representative from 3 independent experiments.
